## Supplementary material for "The virtualome: a computational framework to evaluate microbiome analyses"

### Supplementary Material 1. Ecological structure of microbial populations

A microbial community can be defined as a collection of populations. Populations represent different spatial locations (in the case of soil or water microorganisms for instance) or different host individuals in the case of symbiotic bacteria. Under this definition, microbiomes are populations of microorganisms. The ecological characterization of a community is fully determined by the list of species its contains, together with their abundances. In real-world microbial communities, it has been observed that the abundance of species in a community follows three main macro-ecological rules (defined in [1]). We will briefly summarize these laws. To that end, we will represent a microbial community by a numerical matrix $C$ such that each element, $C_p^s$, represents the abundances of species $s$ in population $p$. We will label as $N_s$ and $N_p$ the number of species and populations in community $C$, respectively.

Let us now define the following variables:

1. *Abundance fluctuation distribution* (AFD): distribution of abundances of a species $s$ across populations:

$$\text{AFD}(s) = \{C_1^s, \cdots, C_{N_p}^s\}$$

2. *Species abundance distribution* (SAD): distribution of species abundances within a given population $p$:

$$\text{SAD}(p) = \{C_p^1, \cdots, C_p^{N_s}\}$$

3. *Mean abundance distribution* (MAD): average abundance of a species $s$ in the populations of the community:

$$\text{MAD}(s) = \frac{C_1^s + \cdots + C_{N_p}^s}{N_p}$$

According to [1], the ecological constrains on species abundance observed in real-world microbial communities translate into the following properties of these variables:

1. The AFD follows a gamma distribution.
2. The SAD and the MAD follow lognormal distributions.
3. The mean and variance of species abundance in the populations of a community exhibit a quadratic relationship (Taylor's law).

The reader is referred to [1] for further details.

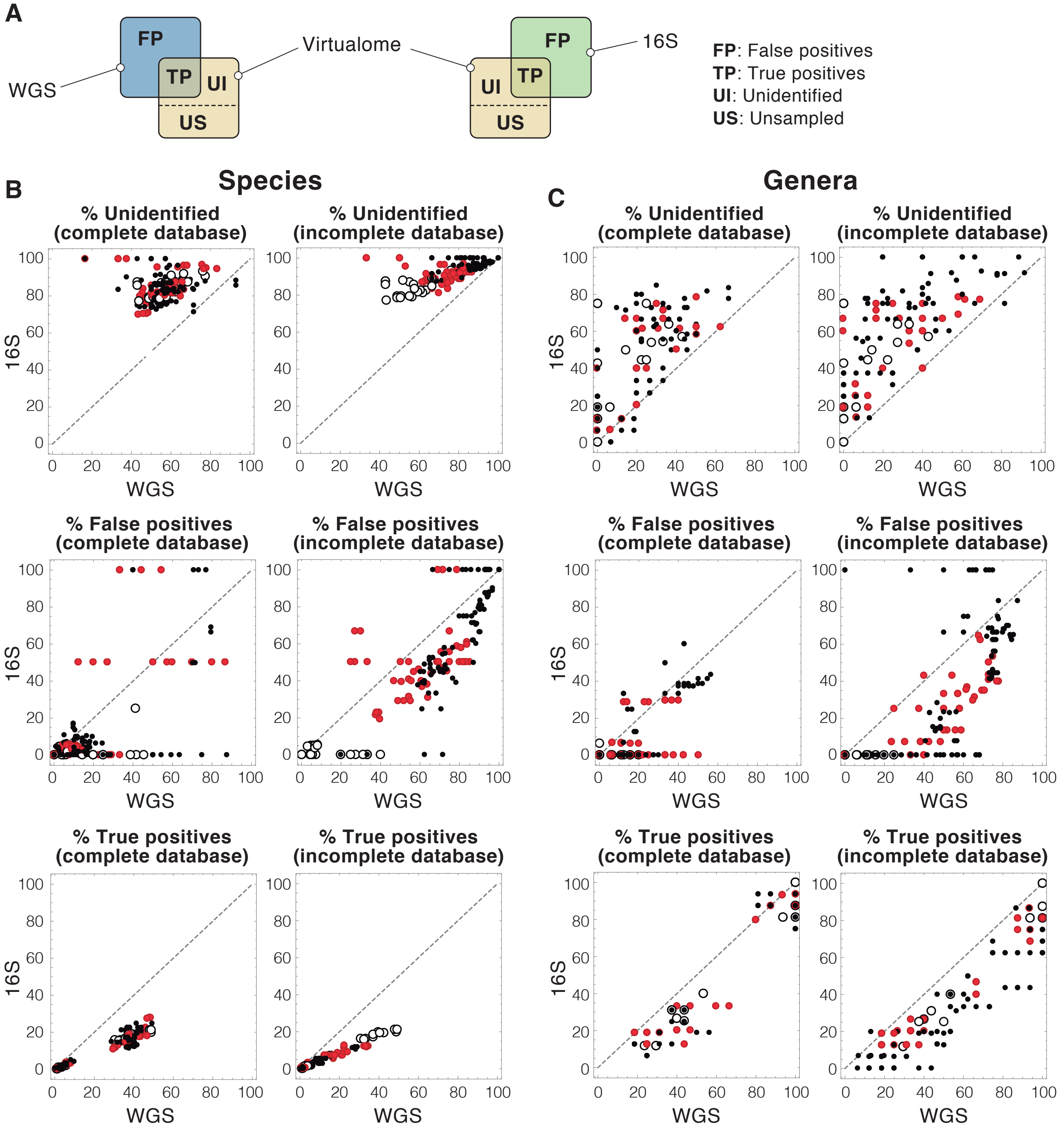


**Figure S1. Comparison between the characterization of virtualomes by 16S and WGS.** A) Venn diagrams showing the logic of the comparison between 16S and WGS. B) Percentage of unidentified species (upper), false positives (medium) and true positives (lower) in 16S vs. WGS analysis of the species composition of virtualomes. C) Same as B at the genus level. (Open dots: 100% of the species or genera of the virtualome are in the databases; Red dots: 50% of species or genera in DBs; Black dots: 25% of species or genera in DBs. Dashed lines: $x = y$.)

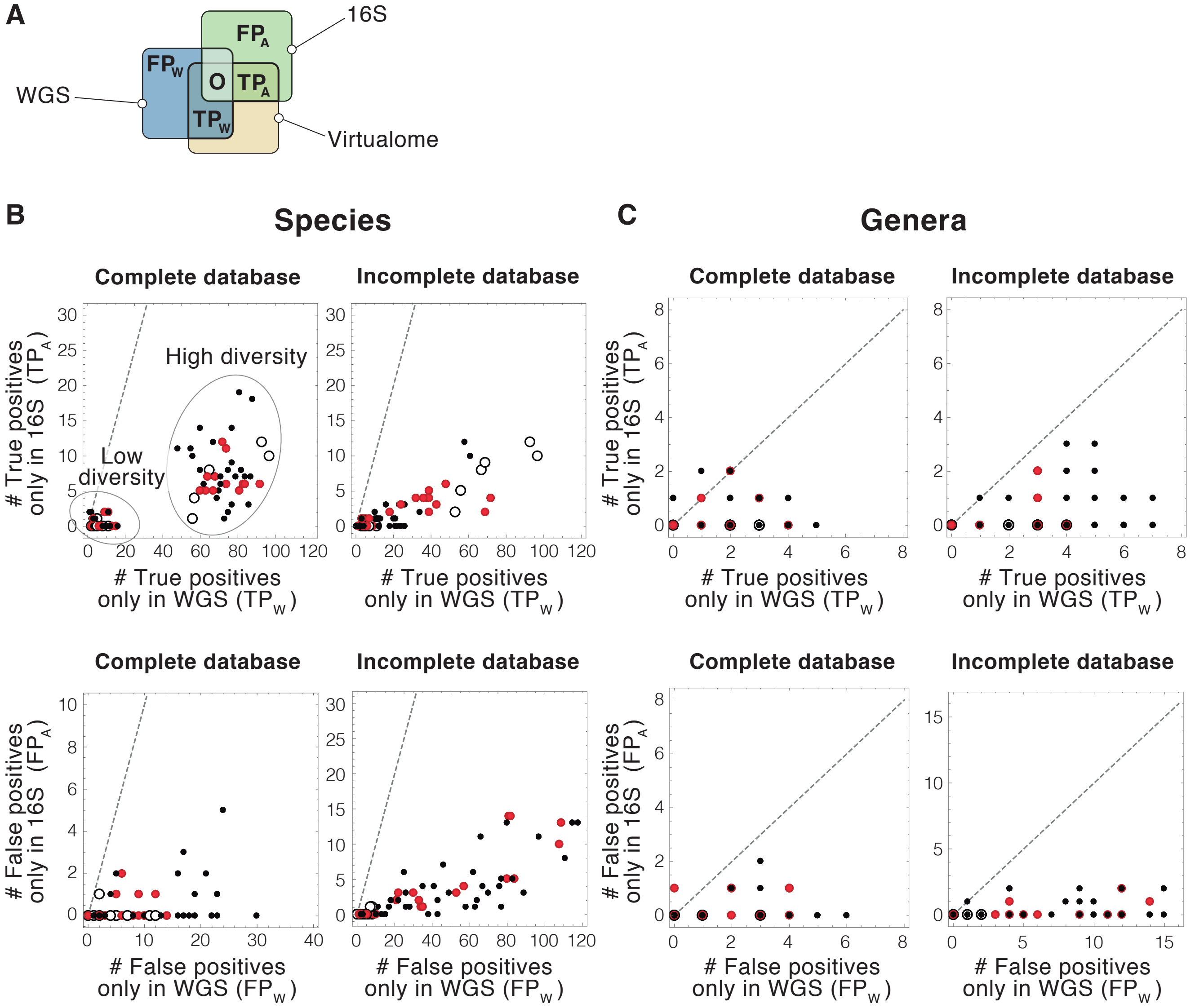


**Figure S2. Differential characterization of virtualomes by 16S and WGS.** A. Venn diagrams showing the sets considered in this figure. O: overlap between 16S and WGS (species and genera simultaneously detected by 16S and WGS; see Fig. 3 in the main text). $FP_W$ and $FP_A$: number of false positives detected only in WGS and 16S respectively. $TP_W$ and $TP_A$: number of true positives detected only in WGS and 16S respectively. B) True positives (upper) and false positives (lower) present exclusively in 16S vs. WGS using complete (left) and incomplete (right) databases. C) Same as B at the genus level. (Open dots: 100% of the species or genera of the virtualome are in the databases; Red dots: 50% of species or genera in DBs; Black dots: 25% of species or genera in DBs. Dashed lines: $x = y$.)

### List of Supplementary Data files

- **Supp. Data 1**: Complete and incomplete databases used in the analyses of virtualomes.
- **Supp. Data 2-4**: Virtualomes analyzed in this work.
- **Supp. Data 5**: Raw data of the amplicon analysis of *G. mellonella* microbiome.
- **Supp. Data 6**: Raw data of the WGS analysis of *G. mellonella* microbiome.
- **Supp. Data 7**: Significant changes of abundance in *G. mellonella* microbiome detected by 16S analysis.
- **Supp. Data 8**: Significant changes of abundance in *G. mellonella* microbiome detected by WGS analysis.

These files can be downloaded from https://data.mendeley.com/datasets/shm68b8x6t/1.
